# Drift-like dynamics and information flow across the cortical hierarchy during working memory

**DOI:** 10.1101/2025.08.07.669201

**Authors:** Hsin-Hung Li, Wei Ji Ma, Clayton E. Curtis

**Affiliations:** Department of Psychology, New York University, New York, NY 10003, USA; Center for Neural Science, New York University, New York, NY 10003, USA; Department of Psychology, The Ohio State University, Columbus, OH 43201, USA

## Abstract

Working memory is supported by widespread and distributed brain regions spanning across the cortical hierarchy. However, how working memory content evolves and is transmitted across cortical regions remains largely unknown. Here, we investigated the flow of working memory information across the cortex using time-resolved fMRI decoding. Across multiple regions in visual and parietal cortex, we found that decoded working memory content drifted over time from the memoranda towards the later errors in memory reports, consistent with drift-like dynamics predicted by attractor models. These behaviorally predictive neural memory errors emerged earliest in higher-order dorsal visual area (V3AB) and intraparietal sulcus (IPS0), and then later in early visual cortex (V1), suggesting a propagation of mnemonic information originated in higher-level visual area. Inter-areal correlation analyses revealed that during memory maintenance, information flowed predominantly in a top-down manner—from higher to lower visual areas whereas during passive viewing feedforward dynamics prevailed. Together, our findings demonstrate that working memory maintenance involves temporally structured drift dynamics and feedback-dominated information flow across cortical hierarchy, providing a mechanistic link between neural population dynamics and the formation of memory errors.

## Introduction

Working memory (working memory) is the cognitive process that holds task-relevant information online over a period of time. working memory is constrained by limited resources, and its precision decreases over delay (Magnussen et al., 1998; McKeown & Mercer, 2012; Pertzov et al., 2017; Phillips, 1974; Shin et al., 2017). Previous studies have conceptualized the maintenance of working memory content and the emergence of its error as a drift diffusion process where the accrual of noise over time perturbs working memory (Fennell & Ratcliff, 2023; Panichello et al., 2019; Schneegans & Bays, 2018). Mechanistically, this process has been modeled using neural networks with attractor dynamics, where memory content is stored in the presence of neural noise (Amit & Brunel, 1997; Bouchacourt & Buschman, 2019; Compte et al., 2000; X. J. Wang, 1999; reviewed in X.-J. Wang, 2021). Findings from single-cell recordings (Finkelstein et al., 2021; Inagaki et al., 2019; Wimmer et al., 2014) and EEG studies in humans (Wolff et al., 2020) are broadly in line with attractor dynamics for working memory. For instance, during working memory delay intervals, population vectors encoding memorized locations from neurons in macaque lateral prefrontal cortex (PFC) drift in the direction of errors in memory-guided saccades generated after the delay (Wimmer et al., 2014); Similarly, neural activity in mouse anterior lateral motor cortex during memory delays converge towards discrete end points that correspond to specific movement directions (Inagaki et al., 2019).

Although the dynamics of working memory have been rigorously characterized in theoretical models and animal neurophysiology, analogous findings from human neuroimaging remain relatively sparse. More critically, because most prior studies focused on single brain regions—particularly the prefrontal cortex—the question of how working memory representations are transmitted across the cortical hierarchy, and how errors in memory evolve across brain regions, remains unknown.

Here, we used time-resolved fMRI decoding to investigate the temporal dynamics of spatial working memory representations across cortical hierarchy. We found that memorized locations can be decoded in all the ROIs along the dorsal visual pathway. We used a memory-guided saccade task like the one used in macaque studies because it yields a continuous metric of memory errors whose direction and amplitude can be modeled. In preview, we found that neural decoding errors drifted from the veridical targets towards errors in the direction of later memory-guided saccades,in line with drift dynamics. Furthermore, by correlating the working memory representations between cortical regions, we found that information flow across cortical regions were task-dependent. Specifically, during memory delays working memory information was dominated by a feedback flow from higher-level visual cortex to early visual cortex

## Results

In a memory-guided saccade task, participants maintained fixation at the screen center and held the location of a briefly presented target in memory over a 12-second retention interval (Figure 1). Target dot was 12° away from the fixation, with its polar angle pseudo-randomly assigned to span an imaginary circle across trials. After the delay, participants reported the target location by generating a memory-guided saccade. Each participant also completed a pRF (population receptive field) session, allowing us to define regions of interest (ROIs) by individual’s visual field maps. Based on prior studies <u>(Harrison and Tong 2009; Rademaker et al. 2019; Sprague et</u> <u>al. 2016; Li et al. 2021; Li and Curtis 2023; Gosseries et al. 2018; Yu and Shim 2017;</u> <u>Hallenbeck et al. 2021)</u>(Gosseries et al., 2018; Harrison & Tong, 2009; Li et al., 2021; Li & Curtis, 2023; Rademaker et al., 2019; Sprague et al., 2016; Q. Yu & Shim, 2017), we focused on the ROIs that exhibited strongest decodable working memory signals. These ROIS include primary visual cortex (V1), extrastriate cortex along the dorsal visual pathway including visual cortex (V2, V3, V3AB) and intraparietal sulcus (IPS0, IPS1, IPS2, IPS3). The current study used data from experiments previously published <u>(Li et al. 2021; Li and Curtis 2023)</u>.

**Figure 1.**
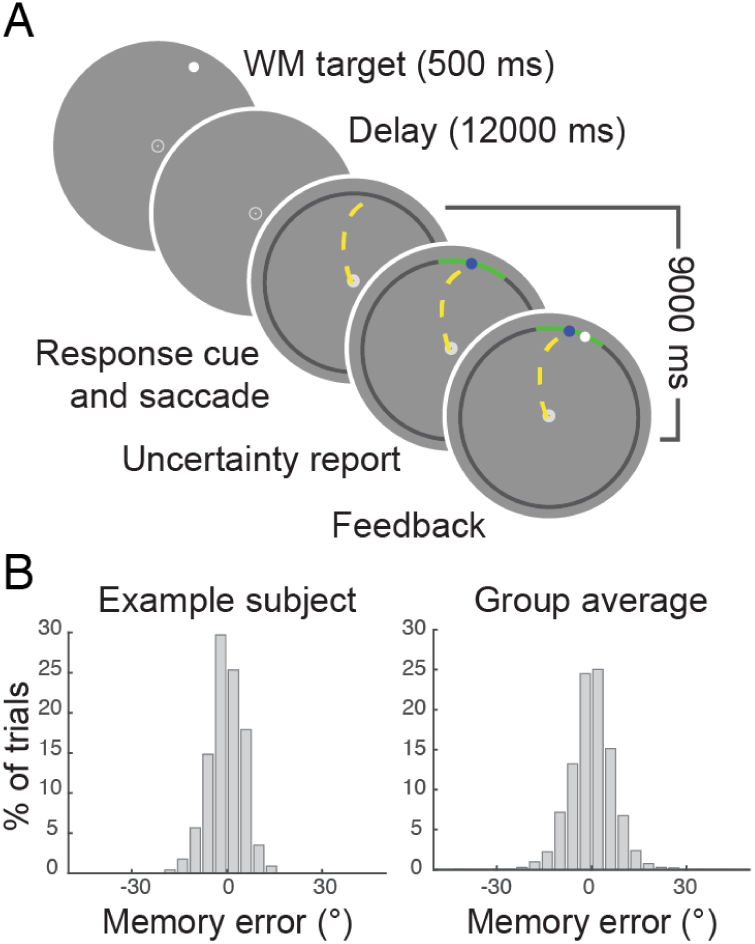
Tasks and behavioral performance. (A) In the memory-guided saccade task, each trial starts with a target presented at 12° eccentricity and a pseudo-randomly assigned polar angle. Participants have to maintain their gaze at the central fixation point throughout the delay, and use a saccadic eye movement to report the memorized location after the onset of the response cue (the onset of the black ring). After the memory report, participants report their uncertainty of their memory by adjusting a confidence interval. Here we focused on the memory reports while the results regarding the uncertainty report were described in a published study (Li et al. 2021). (B) Histograms illustrating the distribution of memory errors in degree polar angle.

To investigate the neural dynamics of working memory, we used a Bayesian method (Li et al., 2021; van Bergen et al., 2015; van Bergen & Jehee, 2021) to decode target locations (polar angles) for single trials using a BOLD signals at each time point during the memory delays (TR = 750 ms). Information about target locations, quantified as the circular correlation between decoded and actual target locations, emerged rapidly in all ROIs (Figure 2A and 2B). In V3 and V3AB, this correlation became significant at about 2.25 seconds (the third TR) into the delay, while decoding in other ROIs emerged at about 3 seconds (the fourth TR). In most ROIs, target information peaked at about 4 to 5 seconds into the delay, consistent with stimulus encoding, and then maintained above-chance throughout the rest of the memory delay. To assess whether these decodable signals were specific to working memory maintenance, we conducted a passive viewing experiment, in which the ‘target’ was a continuous flicker presented throughout the delay, and therefore had no memory demands. Participants were instructed to ignore the flicker while concentrating on an attentionally demanding task at fixation. In this condition, decoding performance also peaked at early time points but then returned to chance levels midway through the delay, indicating that sustained decodability relies on the engagement of the working memory (Supplementary Figure 1).

**Figure 2.**
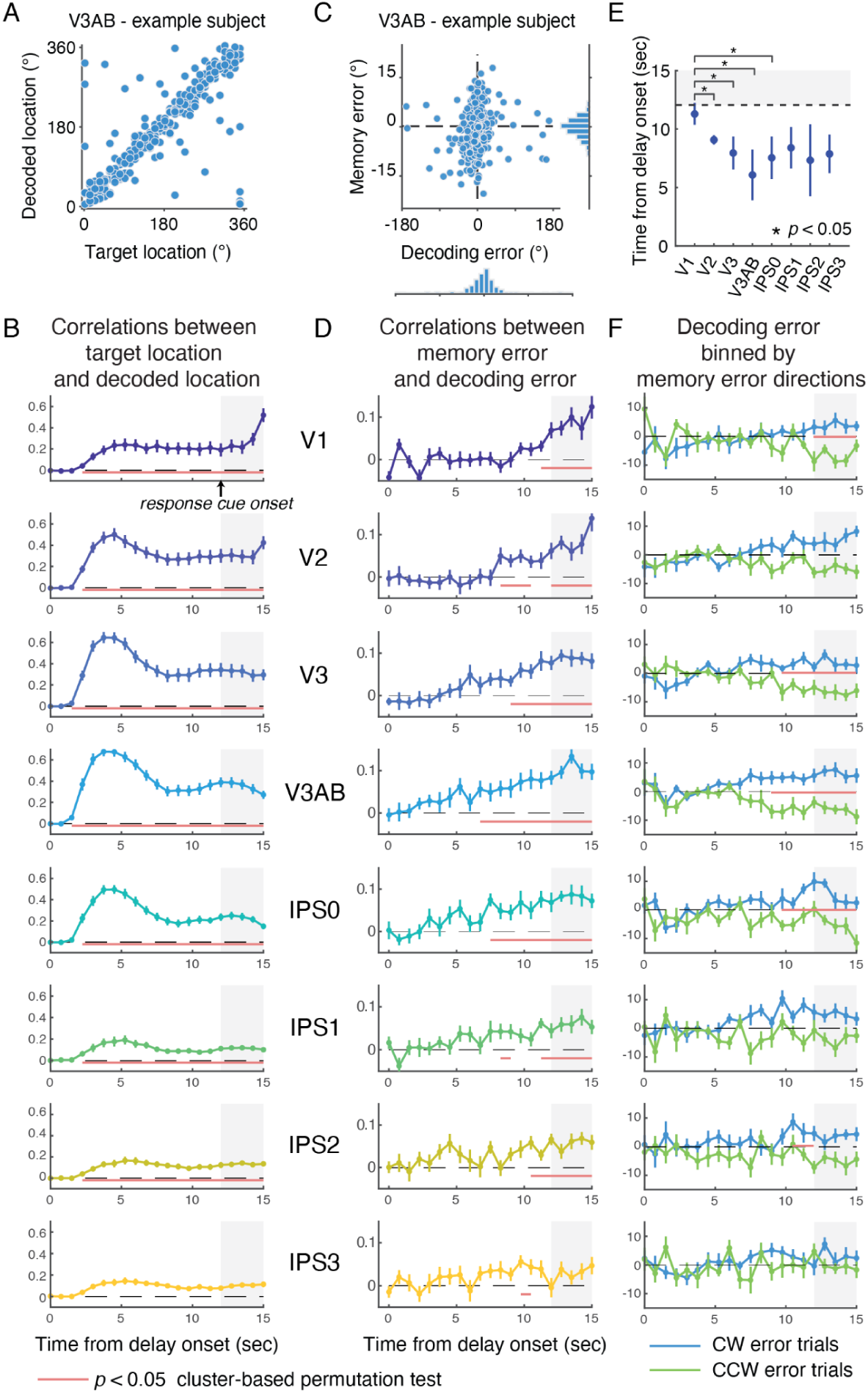
The temporal dynamics of working memory signals. (A) Decoded location and target location from V3AB of an example participant. (B) The time courses of the circular correlations between decoded locations and target locations. The red horizontal lines indicate the clusters where correlations are significantly above zero based on cluster-based permutation tests. The gray shaded area marks the time from the onset of the response cue. The data points represent mean ±1 s.e.m. (C) Decoding error and behavioral memory error of an example participant. (D) The time courses of the circular correlations between decoding errors and behavioral memory errors. (E) The onset times of significant correlations between decoding errors and behavioral memory errors. The error bars represent ±1 bootstrapped standard deviation. (F) The time courses of decoding errors for trials in which the participants made clockwise (blue) and counter-clockwise (green) errors.

We next focused on neural responses that predicted behavioral memory reports beyond the target’s location. We asked whether decoding errors were predictive of behavioral memory errors by correlating the two. We found that the time courses of these error correlations (Figure 2D) diverged markedly from those based on the target locations alone (Figure 2B). In extrastriate visual cortex V2, V3, V3AB and intraparietal sulcus area IPS0, these error correlations gradually ramped up over the delay. To further visualize this drift, we binned trials according to the direction of behavioral memory errors (clockwise CW vs. counterclockwise CCW relative to the target), which confirmed that decoding errors drifted in the direction of upcoming behavioral errors in memory (Figure 2F). We also observed significant error correlations in early visual cortex V1, but the latency of these correlations were significantly delayed relative to those in extrastriate visual cortex and IPS0 (cluster-based permutation test *p* < 0.05, Figure 2D and 2E,). In V1, neural decoding errors were not predictive of behavioral errors in memory until the TR right before the response cue (Figure 2D and 2E). Overall, these error correlations are consistent with the predictions by drift-diffusion processes and neural networks with attractor dynamics (Compte et al., 2000; Wimmer et al., 2014). We observed such dynamics in multiple visual maps across the cortical hierarchy, but with systematic differences in their timing. In the time courses of both Figure 2B and 2D, working memory-related signals in high-level visual area and IPS0 preceded those in early visual cortex, such as V1. This temporal order suggests that working memory content may first be formed in higher-order regions and subsequently propagated backward to early visual cortex. To quantify this information flow during the memory delay, we focused on two areas that showed the earliest and latest drift dynamics—V3AB and V1, respectively.

Next, we computed trial-wise inter-areal correlations of decoded target locations between these two regions (Figure 3A). In Figure 3B, using V1 as the seed region, we correlated its decoded values at each time point with those from V3AB at later time points (a feedforward flow). Conversely in Figure 3C, using V3AB as the seed, we correlated its decoded values with those from V1 at later time points (a feedback flow). Throughout the delay period, inter-areal correlations were consistently stronger when V3AB served as the seed, indicating a predominant flow of information from V3AB to V1 (cluster-based permutation test *p* < 0.05, Figure 3D). We further realigned the inter-areal correlation matrices using V3AB as the reference (Figure 4A) and again found that correlations were stronger when V1 lagged behind V3AB, confirming a dominant feedback flow (Figure 4E). To assess whether this directionality was task-dependent, we applied the same analyses during the passive viewing experiment. In contrast to the working memory task, information flow during passive viewing was stronger in the feedforward direction—from V1 to V3AB—than in the feedback direction (Figure 4C and 4G; also see Figure 3E-G), highlighting that feedback dynamics were specific to working memory. We repeated this analysis using trial-wise decoding errors instead of decoded locations (Figure 4B and 4D). While these correlations were overall weaker, they remained positive—indicating that a portion of the neural variability (or noise) was shared between V3AB and V1, and were detectable in BOLD signals. Crucially, during working memory but not in the passive viewing experiment, the feedback flow from V3AB to V1 remained stronger than the feedforward flow, suggesting that shared noise or variability also traveled predominantly from higher-level visual cortex to early visual areas during memory maintenance (Figure 4F and 4H).

**Figure 3.**
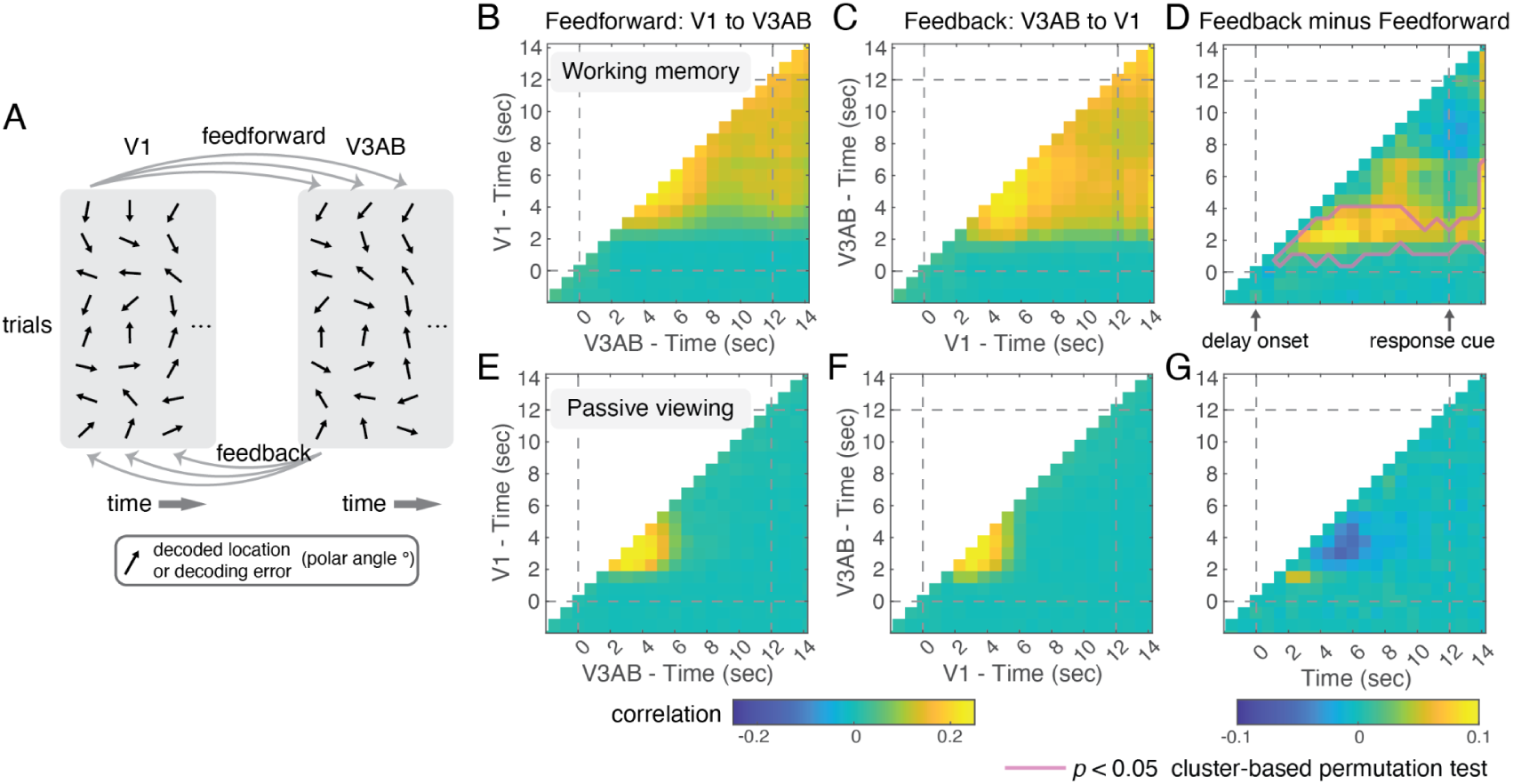
Flow of memory information quantified by inter-areal correlations. (A) Information flows between V1 and V3AB calculated as the correlations of working memory signals (decoded locations or decoding errors) across the two regions and across different time points. Feedforward flow is represented by the correlations between V1’s working memory signals at a given time point and V3AB’s working memory signals at later time points. Feedback flow is represented by the correlations between V3AB’s working memory signals at a given time point and V1’s working memory signals at later time points. (B) Inter-areal correlations based on decoded locations in the feedforward direction. (C) Inter-areal correlations based on decoded locations in the feedback direction. (D) The inter-areal correlations in the feedback direction minus that in the feedforward direction. The pink outline indicates a cluster where the difference is significant based on a cluster-based permutation test. (E) to (G), similar to (B) to (D) but for the passive viewing experiment.

**Figure 4.**
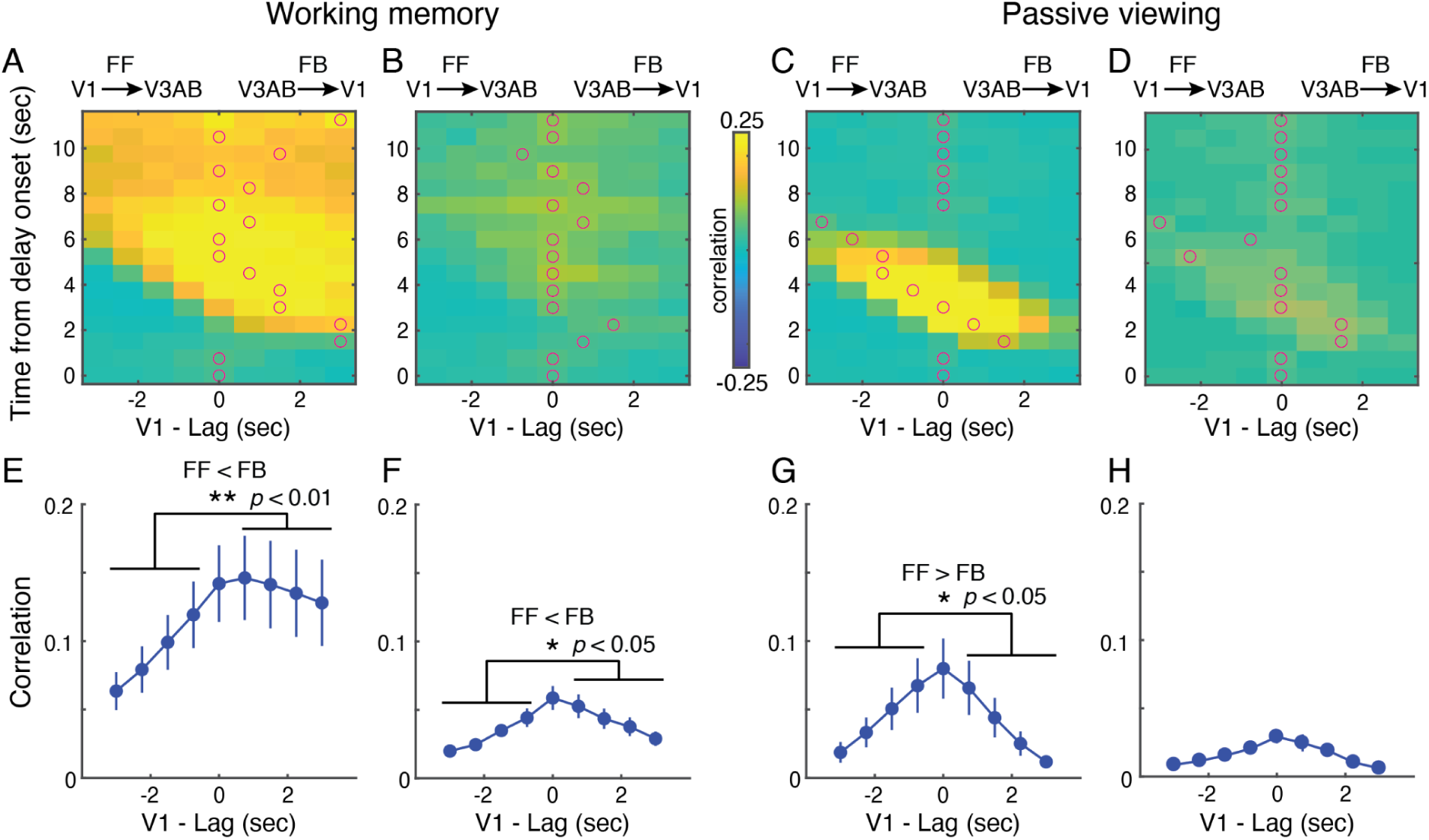
Time lags in inter-areal correlations. (A) Inter-areal correlations based on decoded locations. Similar to Figure 3B and 3C, but the results are realigned so V3AB is the reference time axis (the y-axis), and the decoded locations from V3AB at each time point are correlated with the decoded location from V1 with different time lags (x-axis). Thus, the left half of the image represents feedforward (FF) flow (V1 ahead of V3AB) and the right half of the image represents feedback (FB) flow (V1 lags behind V3AB). In each row, red circles denote peaks in the inter-areal correlations for each reference time point (row). (B) Similar to (A), but here the inter-areal correlations are computed using decoding errors instead of decoded locations. (C) Similar to (A) but for the passive viewing condition. (D) Similar to (B) but for the passive viewing condition. (E) to (H), the averaged inter-areal correlation for each time lag, computed by averaging over each column in the images in (A) to (D) respectively. Paired t-tests were used to compare the overall magnitude of the FF flow (average of the four data points with negative lags) and FB flow (average of the four data points with positive lags). The data points represent mean ±1 s.e.m.

In network models with drift dynamics, memory maintenance is often described as a localized “bump” of activity centered on the memorized target <u>(Wimmer et al. 2014; Compte et al. 2000;</u> <u>Esnaola-Acebes et al. 2022)</u>. To visualize such bump activity, we projected neural responses in V3AB into the coordinates of a two-dimensional visual field, where the center corresponds to the fixation point (see Methods). We visualized the neural activity for three bins of trials sorted by the direction of behavioral memory errors (Figure 5A). Reconstructed neural responses showed a bump at the target’s polar angle (Figure 5B). Comparisons between CW and CCW trials revealed significant differences at locations flanking the target, consistent with two bumps drifting in opposite directions towards the direction of errors (CW or CCW) in the later memory-guided saccades (rightmost panel, Figure 5B).

**Figure 5.**
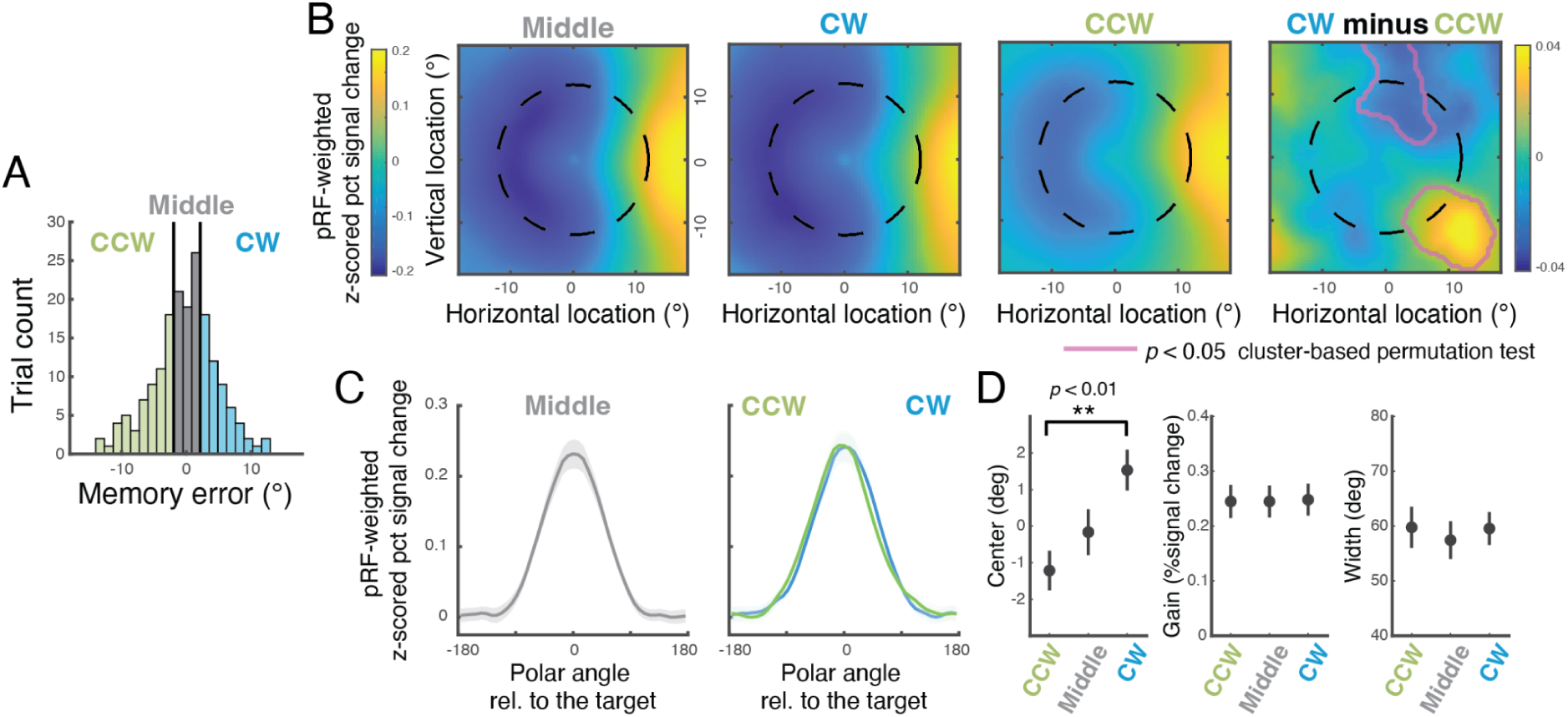
Visualizations of population activity in V3AB. (A) Each participant’s trials were placed into three bins with an equal number of trials based on the direction and magnitude of memory errors. (B) Activation maps visualize the projection of voxel activity patterns onto two-dimensional visual field space. The first three images: Each image represents the activation map reconstructed using the activity of V3AB averaged over the late delay duration (7 to 12 seconds from delay onset). The right-most image, the difference between the activation map of the CW trials and the CCW trials. (C) Polar angle response functions for trials with small errors (‘Middle’ bin, gray curve), CW errors (blue curve), and CCW errors (green curve). (D) Best-fit parameters for polar angle response functions. The data points represent mean ±1 s.e.m.

Previous studies have suggested that memory errors may arise from both a drift of the center of the bump of activity and a decrease in the gain of the bump <u>(Schneegans and Bays 2018;</u> <u>Standage and Paré 2018)</u>. To investigate these two possibilities, we collapsed the two-dimensional reconstructions to one-dimensional response functions over polar angle, visualizing unimodal bumps of activity centered around the polar angle of the targets (Figure 5C). By fitting von-Mises functions to the polar angle response functions, we found that the memory errors were mainly driven by a drift in the centers of the bumps (*p* < 0.01, the main effect of bins in permutation ANOVA; Figure 5D).

## Discussion

We investigated how neural representations of working memory content evolve over time across the cortical hierarchy in humans. We observed drift-like dynamics, with decoding errors systematically drifting toward the direction of subsequent errors in memory, across multiple visual maps in visual and parietal cortex. This error-predictive signal emerged first in V3AB and IPS0 and then later in early visual cortex. Inter-areal correlations of working memory content during working memory maintenance further confirmed that information flowed from high-level visual cortex to early visual cortex. Together, these results provide evidence that working memory maintenance involves drift-like dynamics, not just within, but critically across the cortical hierarchy, and is supported by a feedback cascade from parietal-occipital regions to early visual cortex.

Early studies on the neural mechanisms that support working memory mostly restricted their focus to the lateral PFC (reviewed in Curtis & Sprague, 2021; Funahashi, 2017; Funahashi et al., 1989, 1990). However, accumulating evidence from recent studies has led to the emerging view that representations of working memory content are distributed across multiple brain regions (Ester et al., 2015; Huang et al., 2024; Jerde et al., 2012; Li et al., 2021; Li & Curtis, 2023; Mendoza-Halliday et al., 2014; Panichello & Buschman, 2021; Rademaker et al., 2019; Riggall & Postle, 2012). Whether or how neural representations of working memory may differ and serve different roles across brain regions remains an open question (Christophel et al., 2017; Lee et al., 2013; Panichello & Buschman, 2021; Xu, 2023), but one that deserves attention. Here, we found that among visual field maps along the dorsal visual pathway, all of which exhibited above-chance performance in decoding the working memory target, they showed different temporal dynamics when we analyzed the signals predictive of behavioral memory errors. The results that V3AB and IPS0—the two ROIs sitting at the boundary between the occipital (visual) cortex and the parietal cortex—exhibit the earliest error correlations (Figure 2D and 2E), indicate that they are potentially the source of working memory content. Interestingly, in our previous study, we observed that the neural code of working memory was the most stable in V3AB and IPS0 whereas the neural code of memory targets in V1 underwent significant changes between the stimulus-evoked period to memory maintenance (Li & Curtis, 2023); in another recent study reported that the format of working memory in early visual cortex was ‘aligned’ to that of the IPS during memory delay (Xu, 2023). These previous results and the present findings indicate that the neural representations of the memorized target in early visual cortex are shaped by the working memory content stored in the high-level dorsal visual area and posterior parietal cortex.

This idea that information flows from V3AB to V1 during working memory was corroborated by inter-areal correlations based on decoded location and errors. Previous neurophysiological studies have used regression (Semedo et al., 2019) or canonical correlation (Semedo et al., 2022) to investigate the information flow between cortical regions. Our approach differed in that we first converted multivariate neural responses to the task-relevant dimension—target’s polar angle—and used it to quantify the information flows between pairs of brain regions (Figure 3A). Note that even though the inter-areal correlations computed based on decoded locations were associated with the stimulus locations by definition, they were task-dependent. Indeed, the inter-areal correlations were much stronger during the working memory tasks compared to a passive viewing condition. Moreover, only during working memory did we observe a dominance of feedback flow (Figure 3). The inter-areal correlations computed based on decoding errors were smaller in magnitude compared to those computed using decoded locations, but were still significantly above zero (Figure 4B). These results indicate that while a major proportion of the neural (or measurement) noise was independent between ROIs, some variability may be shared across brain regions. Our analysis showed that such variability shared between brain regions also exhibited stronger feedback flow than feedforward flow during working memory delay.

The present findings echo some previous neuroimaging studies that also reported decoding error or multivariate neural activity to correlate with behavioral errors (Iamshchinina et al., 2021; Li et al., 2021; Wolff et al., 2020). In the literature of perceptual decision-making, this type of choice-related signal, conditioned upon the stimulus, is often considered a strong indicator of a tight link between the measured neural responses and behaviors (Nienborg et al., 2012; Nienborg & Cumming, 2009; Shadlen et al., 1996; X.-J. Yu et al., 2015). Our results here are generally consistent with neurophysiology studies reporting that choice-related or choice probability is weaker in early sensory cortex, and stronger in higher-level regions (Britten et al., 1996; Camarillo et al., 2012; Dodd et al., 2001; Goris et al., 2017; Nienborg et al., 2012; Nienborg & Cumming, 2009; X.-J. Yu et al., 2015). Different from our results, a previous fMRI study reported that the behaviorally-consistent bias in neural decoding was higher in the early visual cortex (EVC) than in IPS regions (Iamshchinina et al., 2021). However, the EVC in this study included combined striate (V1) and multiple extrastriate visual areas (V2 to V4). It is unknown whether the behaviorally-consistent bias they found in EVC was mainly driven by extrastriate cortex like V3. Similarly, their IPS area contained a combined ROI including IPS0 to IPS4. Thus, it remains unclear if our results are inconsistent or not. We found that the error-related signals were early and prominent in IPS0, but were much weaker from IPS1 and beyond. Aggregating all IPS subdivisions into a single ROI may obscure the specific contributions of individual subregions.

When we visualized the neural response of V3AB in the coordinates of the visual field, we observed a bump of activity that drifted in the direction of errors in memory (Figure 5). The drift we observed is predicted by neural networks with drift dynamics (Amit & Brunel, 1997; Bouchacourt & Buschman, 2019; Compte et al., 2000; X. J. Wang, 1999; reviewed in X.-J. Wang, 2021), and resembled results from electrophysiological recordings from macaque PFC while performing a similar memory-guided saccade task (Wimmer et al., 2014). Previous studies have postulated that errors in memory may be caused by drift of the bump of activity or a reduction of the gain <u>(Schneegans and Bays 2018; Standage and Paré 2018)</u>. In our study, focusing on a set size of one, we found that memory errors were primarily attributable to the drift of memory representations. These results are in line with the attractor neural networks where the decay of memory is mainly attributed to the drift of bump activity at the stored feature value <u>(Compte et al. 2000; Wimmer et al. 2014; Burak and Fiete 2012)</u>. However, this does not preclude the possibility that changes in gain contribute to memory variability under different conditions. For instance, in a recent study, we investigated spatial working memory with a set size of two and incorporated spatial attentional cues to manipulate behavioral relevance of the items. We observed that task relevance affected the precision of working memory representations through the change of neural gains (Li et al., 2025). These findings suggest that multiple mechanisms may influence working memory quality. Specifically, both drift and changes in the gain (amplitude) of neural activity can contribute to memory variability, with their relative impact depending on task demands and context.

## Methods

### Subjects

We analyzed the datasets previously reported in Li et al. (2021) and Li & Curtis (2023). There were sixteen participants in total: fourteen were the same participants reported in Experiment 2 in Li et al. (2021), and two additional, non-overlapping participants were from the control experiment reported in Li & Curtis (2023). All participants had normal or corrected-to-normal vision. Written informed consent was obtained prior to participation, following protocols approved by the University Committee on Activities Involving Human Subjects at New York University. Participants were compensated at a rate of $30 per hour.

### Procedures

Participants completed a memory-guided saccade paradigm while undergoing fMRI scanning. Each trial began with the presentation of a working memory (working memory) target—a light gray dot (0.65° in diameter)—for 500 ms. Targets have a fixed eccentricity at 12°, and their polar angle was pseudo-randomly drawn from 32 evenly spaced positions covering a full circle. After the target offset, there was a 12-second delay. During the delay, participants were instructed to maintain fixation at the central fixation point while retaining the spatial location of the target in memory. At the end of the delay, the fixation point changed its appearance (from an empty circle to a filled gray dot) and a black circular ring appeared, centered at fixation and matching the eccentricity of the target. These changes served as the go cue, instructing the participants to indicate the remembered location by executing a saccade to the memorized location. The initial saccade landing location was recorded by the eye tracking, and a dot was shown at that point. Participants could then refine this memory report using a manual dial and confirm their final memory estimate with a button press. Upon confirmation, a visual arc appeared on the ring, centered on the reported location. Participants adjusted the length of this arc using the manual dial to indicate their subjective uncertainty—the longer the arc, the greater the expressed uncertainty. This confidence report was finalized with another button press. Feedback was then provided by displaying a white dot at the actual target location. Participants earned points if the true location fell within the arc boundaries (for task details, see Li et al., 2021). Analyses of the uncertainty reports were presented in a prior publication (Li et al., 2021). Two participants from the control experiment reported in Li & Curtis (2023) completed the same procedure, except that the go cue consisted only of the black ring indicating the target’s eccentricity, without a change in the fixation point.

### Apparatus and Eye Tracking

Visual stimuli were projected using a VPixx ProPix LCD projector positioned behind the MRI bore. Participants viewed the display via an angled mirror with a field of view of 52° horizontally and 31° vertically. A gray circular aperture (30° in diameter) remained visible on the screen throughout the experiment. Eye movements were monitored using an EyeLink 1000 Plus infrared video-based eye tracker (SR Research) inside the scanner bore, with a sampling rate of 500 Hz. During and between scanning runs, gaze data were monitored in real time. The eye tracker is calibrated at the start of each fMRI session, and at the beginning of a run when needed.

### MRI Data Acquisition and preprocessing

MRI data were collected on a Siemens Prisma 3T scanner equipped with a 64-channel head/neck coil. Functional images were acquired with 44 slices and a voxel size of 2.5 mm (4x simultaneous-multi-slice acceleration; FoV was 200 × 200 mm; no in-plane acceleration; TE/TR: 30/750 ms, flip angle: 50 deg, Bandwidth: 2290 Hz/pixel; 0.56 ms echo spacing; P→ A phase encoding). To correct for susceptibility-induced distortions, spin-echo images were intermittently collected during the session in both forward and reverse phase-encoding directions (TE = 45.6 ms, TR = 3537 ms; three volumes per direction) using the same slice prescription but without multiband acceleration. Functional data for retinotopic mapping were acquired in a separate session using a higher-resolution multiband protocol with 56 slices (4x multiband acceleration), a voxel size of 2 mm isotropic, and a FoV of 208 × 208 mm. Imaging parameters for this session were: TR = 1200 ms, TE = 36 ms, flip angle = 66°, bandwidth = 2604 Hz/pixel, echo spacing = 0.51 ms, and phase encoding in the P→A direction. In each participant’s retinotopic mapping session, we also collected 2 or 3 T1 weighted whole-brain anatomical scans (MPRAGE sequence; 0.8 mm^3^).

### Quantification and statistical analysis

#### Behavioral Data Analysis

As participants were allowed to manually refine their saccadic response using a dial, the final adjusted position of the dot was taken as the participant’s memory report. Eye position data were analyzed offline. First, the raw gaze coordinates were smoothed using a Gaussian kernel, then transformed into eye velocity by computing velocity based on a five-point moving window of adjacent samples. Saccades were identified when eye velocity exceeded the median velocity by more than five standard deviations and lasted at least 8 ms. Trials were excluded if the primary saccade could not be reliably detected, if gaze deviated from fixation by more than 2.5°, or if the memory error exceeded three standard deviations.

#### Generative model and Bayesian decoding

We applied a generative model-based approach (Li et al., 2021; van Bergen & Jehee, 2021) to decode the spatial location (polar angle) of the working memory (working memory) targets from the fMRI BOLD signal. The time series data were first converted to percent signal change and then normalized (z-scored) within each run. The decoding procedures followed those described in Li et al. (2021), with one key difference: while the previous study decoded the average BOLD response over a late delay period (i.e., normalized percent signal change averaged from 5.25 to 12.00 seconds after delay onset), here we performed decoding at each individual time point (TR = 750 ms). Below, we summarize the generative model and decoding procedure.

In the generative model, the multivoxel BOLD response associated with a given stimulus location (polar angle) was assumed to follow a multivariate normal distribution. The expected response (mean) of each voxel for a particular stimulus was defined by its tuning curve, representing the voxel’s response profile across polar angles. To estimate each voxel’s tuning, we used a weighted linear combination of eight basis functions that uniformly spanned the polar angle space. These basis functions were raised cosine functions.

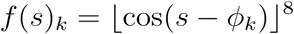

where 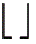 represents half-wave rectification and *ϕ_k_* is the center of the kth channel. The response of ith voxel *b_i_* given a stimulus s is then modeled as

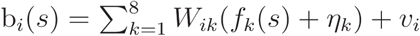

where *W* is a weighting matrix that determines the weights of each basis function for each voxel. Thus, for each training dataset, we assumed that the voxel activity pattern followed a multivariate normal distribution 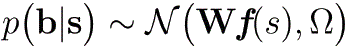, in which 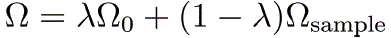.

When the number of variables (voxels) is larger than the number of observations (trials), the sample covariance is not invertible. To ensure a stable estimation of the covariance matrix, the covariance matrix was modeled as the sample covariance matrix Ω_sample_ “shrunk” (Ledoit & Wolf, 2004) to a target covariance matrix—the theoretically important covariance matrix Ω_0_. The free parameter λ determined the degree of shrinkage. Here, the target covariance matrix Ω_0_ is assumed to have a simple structure computed as a weighted sum over a diagonal matrix, a rank-1 covariance matrix, and the covariance depending on the tuning functions *W* of the voxels (see details in (Li et al., 2021; van Bergen & Jehee, 2021)).

For each participant and each region of interest (ROI), we performed decoding of the target’s polar angle using a leave-one-run-out cross-validation approach. In the training phase, the generative model was fit to all trials except those from a single held-out run. The model estimated free parameters based on this training set, and then for each trial in the held-out run (the test set), decoding was performed using Bayes’ rule:

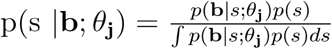

We assumed the prior *p*(*s*) to be uniform, and we approximated the continuous posterior probability function by sampling 1000 steps evenly spanning the location space. Finally, we computed the mean of the decoded posterior probability function to represent the decoded location (polar angle) of each single trial in the testset.

### Statistical tests

To provide robust statistical tests across multiple contiguous time points or spatial locations, we applied cluster-based permutation tests whenever feasible. In Figure 2B, to identify time points where the decoded location correlated with the target location, we computed the circular correlation between the decoded and target locations for each participant. At each time point, we performed a one-sample t-test to evaluate whether the correlation differed from zero. Neighboring time points with uncorrected significance (*p* < 0.05) were grouped into clusters. For each cluster, we summed the t-values of all time points within the cluster. The significance of each cluster was then determined by comparing its summed t-value to a null distribution generated via a permutation procedure. Specifically, we randomly permuted the decoded target locations, repeated the same analysis (t-tests at each time point and clustering of significant points), and computed the maximum cluster-level t-sum for each permutation. This procedure was repeated 1,000 times to generate a null distribution of maximum clusters’ t-sums. Clusters, originally identified in the real data, with t-sums exceeding the 95th percentile of the null distribution were deemed significant and are marked with red horizontal lines in Figure 2B. The same analysis was used for assessing correlations between decoding error and behavioral memory error (error-error correlation; Fig. 2D). To estimate the confidence interval for the onset latency of error-error correlations and compare the latency between ROIs, we performed a bootstrap procedure: participants were resampled with replacement, and for each resampled dataset, the cluster-based test was applied to identify the earliest time point that was deemed significant. This process was repeated 1,000 times, yielding a distribution of onset latencies for each ROI (Fig. 2E). In other analyses where we compared two different conditions (Fig. 2F, 3D, and the rightmost panel of 5B), we used the same cluster-based method, with the null distribution generated by randomly shuffling condition labels across participants. Full details of the cluster-based permutation procedure can be found in Maris and Oostenveld (2007).

---

**Supplementary Figure 1.**
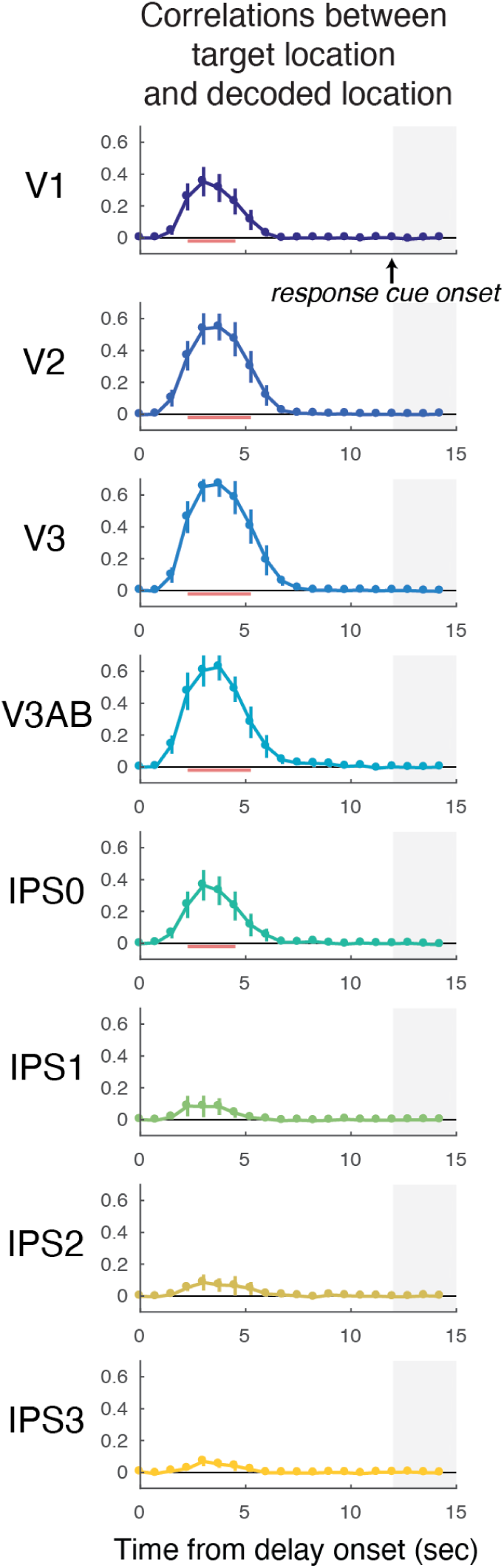
The temporal dynamics of stimulus information in the passive viewing experiment. The figure is plotted in the same format as Figure 2B, but for the passive viewing experiment. The time courses of the circular correlations between decoded locations and target locations. The red horizontal lines indicate the clusters where correlations are significantly above zero based on cluster-based permutation tests. The gray shaded area marks the time from the onset of the response cue.

